# *Schizosaccharomyces pombe* KAT5 contributes to resection and repair of a DNA double strand break

**DOI:** 10.1101/2020.06.03.132316

**Authors:** Tingting Li, Ruben C. Petreaca, Susan L. Forsburg

## Abstract

Chromatin remodeling is essential for effective repair of a DNA double strand break. KAT5 (*S. pombe* Mst1, human TIP60) is a MYST family histone acetyltransferase conserved from yeast to humans that coordinates various DNA damage response activities at a DNA double strand break (DSB), including histone remodeling and activation of the DNA damage checkpoint. In *S. pombe*, mutations in *mst1*^+^ causes sensitivity to DNA damaging drugs. Here we show that Mst1 is recruited to DSBs. Mutation of *mst1*^+^ disrupts recruitment of repair proteins and delays resection. These defects are partially rescued by deletion of *pku70*, which has been previously shown to antagonize repair by homologous recombination. These phenotypes of *mst1* are similar to *pht1-4KR*, a non-acetylatable form of histone variant H2A.Z, which has been proposed to affect resection. Our data suggest that Mst1 functions to direct repair of DSBs towards homologous recombination pathways by modulating resection at the double strand break.

## Introduction

The DNA damage response choreographs multiple cellular activities such as lesion detection, activation of the DNA damage checkpoint, break repair, and recovery (rev. in (Langerak and Russell 2011)). Chromatin remodeling is central to the faithful execution of these processes, requiring the coordination of histone modifiers, remodelers, and chromatin assembly factors (rev. in (Bakkenist and Kastan 2015; Clouaire and Legube 2019; Aleksandrov *et al*. 2020)). Acetylation of histone H4 is one marker associated with chromatin remodeling at a double strand break (DSB), a potentially lethal form of DNA damage. There is good evidence that histone H4 acetylation is linked to radioresistance and the repair response, possibly by facilitating chromatin mobilization (rev. in (Dhar *et al*. 2017; Van and Santos 2018)).

The MYST family of histone acetyltransferases (HATs) is defined by a distinct catalytic domain, and its members are implicated in histone acetylation at DSBs (rev. in (Utley and Côté 2003; Thomas and Voss 2007)). KAT5 is the most highly conserved HAT of this family, known as Mst1 in *S. pombe*, Esa1 in *S. cerevisiae*, and TIP60 in mammals (rev. in (Avvakumov and Côté 2007; Pillus 2008)). The gene is essential for viability in all species, because in addition to its role in DSB repair it performs other functions such as transcriptional regulation, heterochromatin assembly and centromere assembly (rev. in (Pillus 2008)). In humans, TIP60 has also been identified as a tumor suppressor (rev. in (Sapountzi *et al*. 2006; Voss and Thomas 2009) and is a therapeutic target (rev. in (Judes *et al*. 2015)).

KAT5 contains a chromodomain in addition to its HAT domain, which suggests that it binds methylated histones (rev. in (Utley and Côté 2003; De La Cruz *et al*. 2005)). KAT5 functions in the context of a large protein complex called NuA4, and acetylates numerous substrates including histones H2A, H3, and H4, and the histone variants H2A.X and H2A.Z (Kimura and Horikoshi 1998; Smith *et al*. 1998; Allard *et al*. 1999; Galarneau *et al*. 2000; Babiarz *et al*. 2006; Keogh *et al*. 2006). Evidence from human cells has shown that KAT5 can acetylate non-histone proteins as well, including the checkpoint kinase ATM and the tumor suppressor TP53 (Sun *et al*. 2005; Jiang *et al*. 2006; Leduc *et al*. 2006; Sykes *et al*. 2006; Tang *et al*. 2006; Kim *et al*. 2007; Naidu *et al*. 2012; Ortega-Atienza *et al*. 2016). Importantly, NuA4 cooperates with the Swr1 chromatin remodeler that is required for exchange of histone H2A with the variant H2A.Z ((rev. in (Scacchetti and Becker 2021) (Kobor *et al*. 2004; Krogan *et al*. 2004; Keogh *et al*. 2006; Zhou *et al*. 2010)).

Evidence from both yeast and humans shows that the NuA4 complex is recruited to DSBs by multiple binding partners including phosphorylated H2A(X) (Downs *et al*. 2004), methylated histone H4K20 (Jacquet *et al*. 2016), methylated histone H3K36 (Li and Wang 2017), methylated histone H3K9 via the KAT5 chromodomain (Sun et al. 2005; Sun et al. 2009, Ayrapetov et al. 2014), and interaction with Nbs1, a component of the end-binding resection complex MRN (Cheng *et al*. 2018). In fission yeast, physical association between KAT5/SpMst1 and the homologous recombination protein Rad52 has also been demonstrated (Gómez *et al*. 2008). However, how these recruiting partners cooperate or compete with each other is unknown.

Recruitment of the NuA4 complex to the sites of DSBs has several outcomes. There is evidence that the recruitment of variant H2A.Z and its subsequent acetylation by KAT5 are an essential early step in the DNA damage response (Morillo-huesca *et al*. 2010; Papamichos-Chronakis *et al*. 2011; Xu *et al*. 2012; Gursoy-Yuzugullu *et al*. 2015). H2A.Z must be acetylated and then removed so that KAT5 can acetylate H4 (Bird *et al*. 2002; Downs *et al*. 2004; Doyon *et al*. 2004; Murr *et al*. 2005; Clarke *et al*. 2017). Acetylation of the damage specific phosophorylated histone variant γ-H2A(X) promotes its turnover (Ikura *et al*. 2007; Jha *et al*. 2008; Sharma *et al*. 2010; Soria *et al*. 2012). There is also evidence that NuA4 acetylates the ssDNA binding protein RPA, to regulate resection (Cheng et al. 2018; Kobayashi et al. 2010; Xu et al. 2010). This activity may be important to activate the homologous recombination pathway as opposed to other types of repair (Renaud *et al*. 2015; Jacquet *et al*. 2016).

In fission yeast, Mst1 has multiple activities in chromosome structure, heterochromatin assembly, centromere function, and transcription (Minoda *et al*. 2005; Gómez *et al*. 2008; Kim *et al*. 2009; Nugent *et al*. 2010; Xhemalce and Kouzarides 2010). Previously, we showed that the temperature sensitive mutation *mst1*^*ts*^ (*mst1Δ nmt:mst1-L344S*) results in sensitivity to DNA damaging agents even at permissive temperatures (Gómez et al. 2008; Gómez, Espinosa, and Forsburg 2005). Although we showed that *mst1*^+^ affects transcription of a variety of genes, changes in expression do not appear sufficient to account for its DNA damage sensitivity (Nugent *et al*. 2010). Here, we present evidence that *mst1*^+^ affects DSB repair in fission yeast. We show that Mst1 is recruited to DSB. *mst1-L344S* cells show defects in recruiting downstream factors including Rad52 and RPA. Interestingly, this phenotype is partially rescued by deletion of the end-binding protein Pku70, normally required for NHEJ repair. Our data suggest that Mst1 contributes to efficient resection. Finally, a non-acetylatable H2A.Z mutant (*pht1-4KR*) phenocopies *mst1-L344S*, indicating that H2A.Z may be an important substrate for this effect.

## Materials and methods

### Strains and Media

Fission yeast cells were grown in YES (yeast extract with supplements) or PMG (pombe minimal glutamate) with appropriate supplements (Sabatinos and Forsburg 2010). Yeast strains used in this research are listed in Supplementary table S2 in the supplemental material.

### Serial Dilution plating

Yeast cell cultures were grown at 25°C in YES for two days. Cultures were diluted in YES to equal concentrations. Five-fold serial dilutions of the cultures were then spotted onto YES plates containing different concentrations of drugs. Plates were incubated at 25°C or 32°C for the time indicated in each figure legend.

### Determination of DSB-induced recombination rates

DSB-induced recombination outcomes were determined using the strain *ade6-M26 int::pUC8/ura4+/MATa/ade6-L469* described in (Schuchert and Kohli 1988; Fortunato *et al*. 1996). The *S. cerevisiae* homothallic endonuclease (HO) was expressed on a plasmid from the *nmt1* promoter. Cells were first grown up in PMG (pombe minimal glutamate) liquid media lacking uracil and thiamine (PMG -ura -thia) for two days. Cell cultures were then plated onto PMG -ura -leu +thia agar media for five days at 25°C. After the five-day incubation, colonies were then streaked onto PMG -leu -thia agar media and incubated at 25°C for seven days (HO expression induced). Three colonies from each induced-break plate were then selected and serially diluted 1:100 in sterile MilliQ H2O. 100μl of the diluted cultures was plated onto PMG low ade -leu - thia agar media. Agar plates were incubated at 25°C. After the 14-day incubation, red, white and red/white half sectored colonies were counted and marked. Plates were then replica plated onto PMG -leu -ade -thia and PMG -ade, -ura -thia and allowed to grow for another six days at 25°C before counting colonies with each phenotype. At least three biological replicates were performed. About 8% are likely “complex” rearrangement, for example colonies that were red/white half sectored on PMG low ade -leu -thia agar media. These colonies arise from recombination intermediates that were not repaired before replication (Fortunato *et al*. 1996).

### Live-cell imaging and Quantitative Measurements

Live cells imaging was performed as described in (Green *et al*. 2015). Yeast cell cultures were grown at 25°C in YES overnight. Cells were transferred into PMG + HULAA (Histidine, Uracil, Leucine, Adenine, Arginine) liquid cultures at 25°C for 16 hours before treating with the indicated drugs for four hours. Cells were collected at OD_595_ 0.3 −0.6, concentrated by centrifuging 1ml at 1500g for 1min and resuspended in 40μl. Concentrated cells were placed on a thin-film pad of 2% agarose in PMG+HULAA on a glass slide. A coverslip was added. Live cells were imaged on DeltaVision Core microscope with softWoRx v4.1 (GE, Issaquah, WA), using a 60X lens, and then deconvolved and projected in softWoRx software. Images were acquired in 13 0.2µm z-sections, then deconvolved and Maximum Intensity Projected (softWoRx, default settings). Two separate fields were imaged in each experiment, and 2 to 4 biological replicates were assessed for consistency. Images for publication were contrast adjusted using an equivalent histogram stretch on all samples. Color balance was adjusted, and scale bars were added in Fiji (Schindelin *et al*. 2012). Significance was calculated using the Mann-Whitney U test.

### LacO LacI-mCherry DSB array colocalization with CFP-tagged Rad52

Co-localization was performed as described in (Yu *et al*. 2013). Cells were cultured at 25°C in PMG + HULAA (Histidine, Uracil, Leucine, Adenine, Arginine) +Thiamine liquid media to OD_595_ of 0.4-0.6. Cells were then washed twice with PMG +HULAA medium and incubated at 25°C for 19 hours to induce HO-driven DSB break. Following induction, cells were collected at OD_595_ of 0.3 to 0.6, and were processed and imaged as described above.

### Chromatin Immunoprecipitation (ChIP) assay

Chromatin immunoprecipitation (ChIP) was performed as in (Du *et al*. 2006). Cells were grown in PMG+HULAA+Thiamine overnight to OD_595_∼0.6. Cells were washed 3 times with EMM+HULAA and released in PMG+HULAA-Thiamine at 32°C with shaking. Samples were taken after washing into PMG+HULAA-Thiamine (T=0), 22hrs(T=22), 24hrs (T=24) and 26hrs (T=26). All samples were fixed with 1% formaldehyde for 20min followed by 5 minutes quenching with Glycine. Cells were lysed in standard ChIP buffer and immunoprecipitations were set up using antibodies against GFP (Abcam290), V5(Abcam15828). DNA was column purified using a Qiagen PCR purification kit and qPCR was performed. Fold enrichment was calculated as 100*^(Adjusted 100% INPUT – Ct(IP)).

### Western Blots

For the time course experiment, cells were grown at 32°C in YES to OD_595_∼0.6 then treated with 40μM camptothecin (CPT) for 4hrs. Cells were washed three times with YES, then released in YES without drug. Time points were taken at T=0 (before addition of drug), T=4 (after 4hrs of drug treatment), and T=6, 8 (two and four hours after removal of drug). Cells were lysed in 20% TCA. Antibodies against phosphorylated H2A (Abcam 17353) and control PCNA (Santa Cruz-56 PC10) were used. For the Chk1-HA phosphorylation experiment, cells were grown at 32°C in YES to OD_595_∼0.6 then treated for 4hrs with 40µM camptothecin, 15mM HU or 0.01%MMS then cells were processed as described above. An anti-HA antibody (12CA5 Abcam16918) was used.

### Reagent, software, and data availability

All strains and other reagents are available to other investigators upon request. All data are provided within the figures and manuscript or in Supplemental Data uploaded to the GSA Figshare portal. Fig S1: *mst1-L344S* sensitivity to phleomycin. S2: Examples of repair protein focus formation in wild type and mutants. S3: *pku70Δ* does not affect cell length. S4: Genetic interactions between *mst1-L344S* and checkpoint mutants. Table S1: Median and quantiles of data in Fig 2A and B. S2: Strain list.

## Results

### Mst1 is important for double-strand break repair in *S. pombe*

Mst1 homologs in mammalian cells (TIP60) and *S. cerevisiae* (*ESA1*) are known to be important for DSB repair (rev. in (Ghobashi and Kamel 2018; Van and Santos 2018; Clouaire and Legube 2019; Aleksandrov *et al*. 2020)). We have previously shown that a *S. pombe mst1*^+^ temperature sensitive allele (*mst1-L344S*) renders cells sensitive to various DNA damaging drugs (Gómez *et al*. 2008). Here we expand these studies to further dissect the role of *mst1*^+^ in DSB repair.

Our original *mst1-L344S* construct is expressed from the *nmt1* promoter inserted at the *leu1*^+^ locus in a deletion background (*mst1Δ nmt:mst1-L344S* Gómez et al. 2008) but subsequently, the same allele was engineered at the native locus under control of the endogenous *mst1*^+^ promoter (*mst1-L344S* Garabedian et al. 2012). We tested the sensitivity of both versions to bleomycin, which creates single and double strand DNA breaks (Stubbe and Kozarich 1987; Burger 1998). Both these strains were modestly sensitive to the drugs at 25°C (Figure 1A; Gómez et al. 2008), consistent with a requirement for Mst1 in DSB repair. Because the *mst1-L344S* allele does not generate autofluorescence to the same extent as our original allele, we employed it here for visual assays.

**Figure 1.**
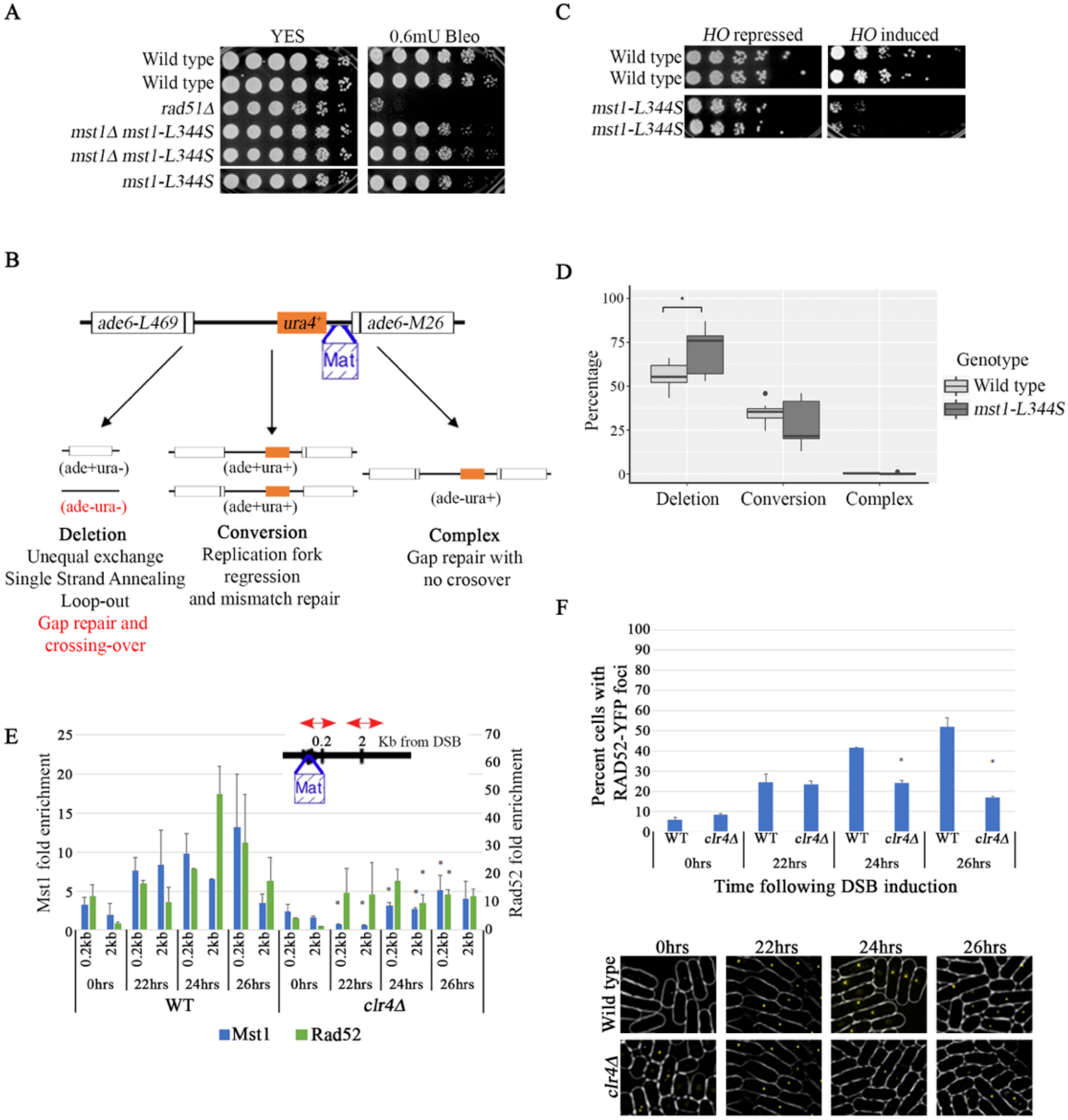
The *mst1-L344S* allele affects DSB repair. **A**. *mst1-L344S* are sensitive to bleomycin. Cells were grown in YES media overnight at 25°C then 5X serial dilutions were spotted onto YES plates or YES plates containing bleomycin. Plates were incubated at 25°C and photographed after 4 days. 1 unit of bleomycin = 1 mg (by potency). **B**. An assay to test chromosomal instability from (Osman *et al*. 1996). Two non-functional *ade6* alleles flank the functional *ura4*^+^ gene. *the ade6-L469* has a mutation at the C terminus while the *ade6-M26* allele has a mutation at the N-terminus (indicated by the vertical lines). Several recombination outcomes are possible and are described in results and methods. **C**. Cells were grown in YES media overnight at 25°C then 5X serial dilutions were spotted onto YES (*HO* repressed) or PMG without thiamine (*HO* induced). Plates were incubated at 25°C and photographed after 5 days. **D**. Percent of deletion, conversion and complex rearrangements after double strand break induction in WT and *mst1-L344S* using the assay in **B**. The bold line in each box represents median and the filled circles represent outliers. Asterisks indicate that recombination percentage is significantly different than WT (p<0.05, Mann-Whitney Test). **E**. Chromatin immunoprecipitation assay to test recruitment of Rad52-YFP and Mst1-V5 to a single DNA double strand break in WT and *clr4Δ*. Using a construct from Du, Nakamura, and Russell 2006 with one single double strand break can be made with the *HO* endonuclease and protein interaction with chromatin can be monitored at 0.2 and 2KB from the break. Following break induction, time points were taken at 0, 22, 24 and 26 hours. qPCR was done with primers provided in Du, Nakamura, and Russell 2006. Asterisks represent fold enrichment is significantly different than that in WT (p<0.1, Student’s t-test). **F**. *clr4Δ* had lower percentage of cells with Rad52 foci after single DSB induction. **Top:** Percentage of cells with Rad52 foci using a construct from Du, Nakamura, and Russell 2006 with one single double strand break can be made with the *HO* endonuclease. Following break induction, cultures were taken at 0, 22, 24 and 26 hours as in Figure 1E and cells were imaged under microscope. Asterisks represent fold enrichment is significantly different than that in WT (p<0.05, Student’s t-test). **Bottom:** Examples of WT and *clr4Δ* cells with Rad52 foci upon DSB induction.

We investigated the role of *mst1*^+^ in repair of a single DSB break. We first utilized a strain that uses the homothallic switching endonuclease (HO) to induce a single DSB between two non-tandem repeats as originally described by (Schuchert and Kohli 1988) and modified by (Osman *et al*. 2000) (Figure 1B). In this strain, a functional *ura4*^+^ gene is flanked by two full length non-functional *ade6* alleles, each containing a different point mutation. The HO endonuclease makes a DSB at *MATa* site adjacent to *ura4*^+^. The repair outcomes can be determined by selecting Ade^+^ cells and screening for Ura^+^ (Fortunato *et al*. 1996; Osman *et al*. 1996, 2000; Ahn *et al*. 2005). Uracil auxotrophic colonies (“deletion type”, whether Ade^+^ or Ade^-^) result from deletion and rearrangement, in which cells repaired the lesion by <u>S</u>ingle <u>S</u>trand <u>A</u>nnealing (SSA), <u>N</u>on-<u>H</u>omologous <u>E</u>nd-<u>J</u>oining (NHEJ), or more complex rearrangements. In contrast, Ade^+^ colonies that retain Ura+ (“conversion type”) are presumed to derive from gene conversion via homologous recombination.

Following induction of the DSB, we observed that the survival of *mst1-L344S* was much lower than the survival in wild-type cells (Figure 1C), consistent with a repair defect. In wild-type cells, about 55.64% of colonies are deletion type (Ade+/Ade-Ura-) while 36.70% of colonies were conversion type (Ade+ Ura+) (Figure 1D). In the surviving *mst1-L344S* cells, we observed a significantly higher percentage of colonies showing deletion-type repair (75.3%) than in wild type (Figure 1D). These data suggest that wild type *mst1*^+^ biases repair towards conversion-type, error-free recombination.

To examine whether Mst1 physically localizes to DSB breaks as has been reported for *Hs*TIP60 and *Sc*Esa1 (Downs and Jackson 2004; Sun *et al*. 2009), we employed a strain previously engineered to test recruitment of proteins by ChIP to one single double strand break (Figure 1E) (Du *et al*. 2006). This assay can monitor both temporal and spatial recruitment of proteins to a single DSB. We detected Mst1 localized near the DSB after induction, although the level of enrichment was lower than that of Rad52 (Figure 1E, note two different Y-axes for Mst1 and Rad52). As expected, Mst1 localization to sequences near DSB was reduced in cells missing H3K9 methyltransferase Clr4 (Figure 1E), which is consistent with previous finding that *Hs*TIP60 binds to the DSB via H3K9^me^ (Sun et al. 2005; Sun et al. 2009, Ayrapetov et al. 2014). Additionally, Rad52 localization to the DSB was reduced in *clr4Δ* (Figure 1E and F), consistent with findings in human cells that H3K9^me^-deficient cells had reduced homologous recombination-mediated repair (Ayrapetov *et al*. 2014, Sun et al. 2009).

### Mst1 contributes to recruitment of repair factors to DSBs

Consistent with a role for Mst1 in DSB repair, we found that the mutant *mst1* at permissive temperature is modestly sensitive to agents that induce DSBs, including the radiomimetic drugs Bleomycin and its analogue Phleomycin (Belenguer et al. 1995) (Figure 1A, supplementary figure 1), suggesting this mutant has attenuated function even under permissive conditions. We next examined the recruitment of homologous recombination proteins in *mst1-L344S* cells using a visual screen following treatment with phleomycin. First, we looked at the localization of RPA and Rad52 proteins (Figure 2A, supplementary table 1). Untreated wild type and *mst1* cells had a similar percentage of cells with foci. However, following phleomycin treatment, we observed significantly more cells with foci in wild type than in *mst1-L344S* cells. We also imaged cells with GFP-tagged Rad54, a recombination protein that acts downstream of Rad52 (Figure 2B; Onaka et al. 2016). Similar to RPA and Rad52, the fraction of *mst1-L344S* cells with Rad54 foci was lower than in the wild type strain (Figure 2B, supplementary table 1). These observations suggest that Mst1 is required for the normal recruitment of downstream HR factors.

Next, we examined the recruitment of Rad52 at a single, induced DSB site. We utilized a strain that contains an HO-inducible DSB adjacent to a lacO array that recruits mCherry-lacI, and CFP-tagged Rad52 (Yu *et al*. 2013). We imaged both wild-type and *mst1-L344S* cells over a four-hour period. In wild-type cells, we observed that 75% of total Rad52 foci co-localize with mCherry signal until 100 minutes (Figure 2C top). After 100 minutes, the percentage of colocalization dropped in wild type, because the mCherry-LacI signal disappears (Figure 2C bottom and Figure 2D). This loss of mCherry-lacI likely reflects resection at DSBs that eliminates the LacO target, as shown previously (Yu *et al*. 2013; Leland *et al*. 2018). Consistent with these prior reports, we observed that the mCherry-DSB foci dropped to 25% by the end of four-hour time-course (Figure 2D, Yu et al. 2013). In contrast, in *mst1-L344S* cells, fewer than 40% of total Rad52 foci colocalize with mCherry-DSB foci across the four-hour time-course, suggesting Mst1 is involved in Rad52 recruitment (Figure 2C top). Strikingly, however, over 90% of mCherry-DSB signal remained even at the end of the time-course (Figure 2C bottom and Figure 2D). This suggests that in the absence of functional Mst1, resection is inhibited, so the LacI-mcherry bound to the lacO array persists.

**Figure 2.**
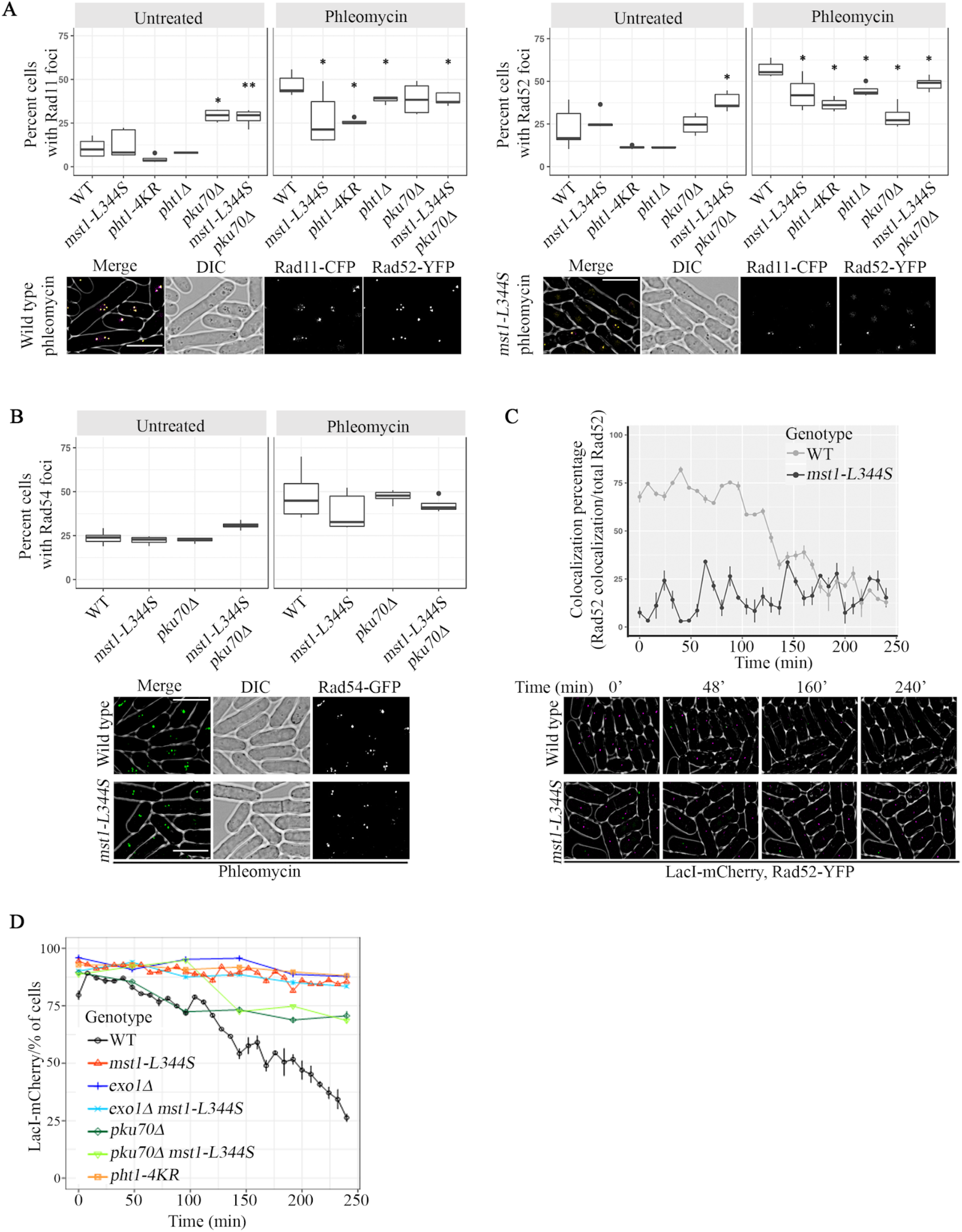
The *mst1-L344S* mutant affects recruitment of repair proteins at 25°C. **A. Top:** Percentage of cells with RPA (Rad11) and Rad52 foci in WT, *mst1-L344S, pht1-4KR, pht1Δ, ku70Δ* and *ku70Δ mst1-L344S*. The bold line in each box represents median and the filled circles represent outliers. Asterisks represent percentage of cells with foci is significantly different than that in WT (*: p<0.05, **: p<0.01, Mann-Whitney Test) **Bottom:** Examples of WT and *mst1-L344S* cells with Rad11 and Rad52 foci. Scale bar is 10µm. Rad11-CFP is colored magenta for visibility. **B. Top:** Percentage of cells with Rad54 foci in WT and *mst1-L344S, pku70Δ and pku70Δ mst1-L344S*. The bold line in each box represents median and the filled circles represent outliers. **Bottom:** Examples of WT and *mst1-L344S* cells with Rad54 foci under phleomycin treatment. Scale bar is 10µm. **C. Top:** Percentage of cells with colocalized Rad52-CFP and HO-induced mCherry foci over 240 minutes. **Bottom:** Representatives of WT and *mst1-L344S* cells with Rad52-CFP and HO-induced mCherry foci over time. Rad52-CFP is colored green for visibility. **D**. Percentage of cells with mCherry-LacI signal over 240 minutes. Image was taken every 8 min in wild-type and *mst1-L344S*, and every 48 min in other mutants.

### Processing of DSBs and a role for Ku

The heterodimeric Ku complex is very abundant and binds rapidly and immediately to DSBs (Fell and Schild-Poulter 2015; Shibata *et al*. 2018). Ku inhibits initial resection driven by MRN, while MRN removes Ku to promote proper resection and facilitate homologous recombination (HR) (Tomita *et al*. 2003; Langerak *et al*. 2011; Shao *et al*. 2012; Myler *et al*. 2017; Shibata *et al*. 2018). Unexpectedly, we found that *pku70Δ* partially rescues the bleomycin sensitivity of *mst1Δ mst1-L344S* (Figure 3A).

**Figure 3.**
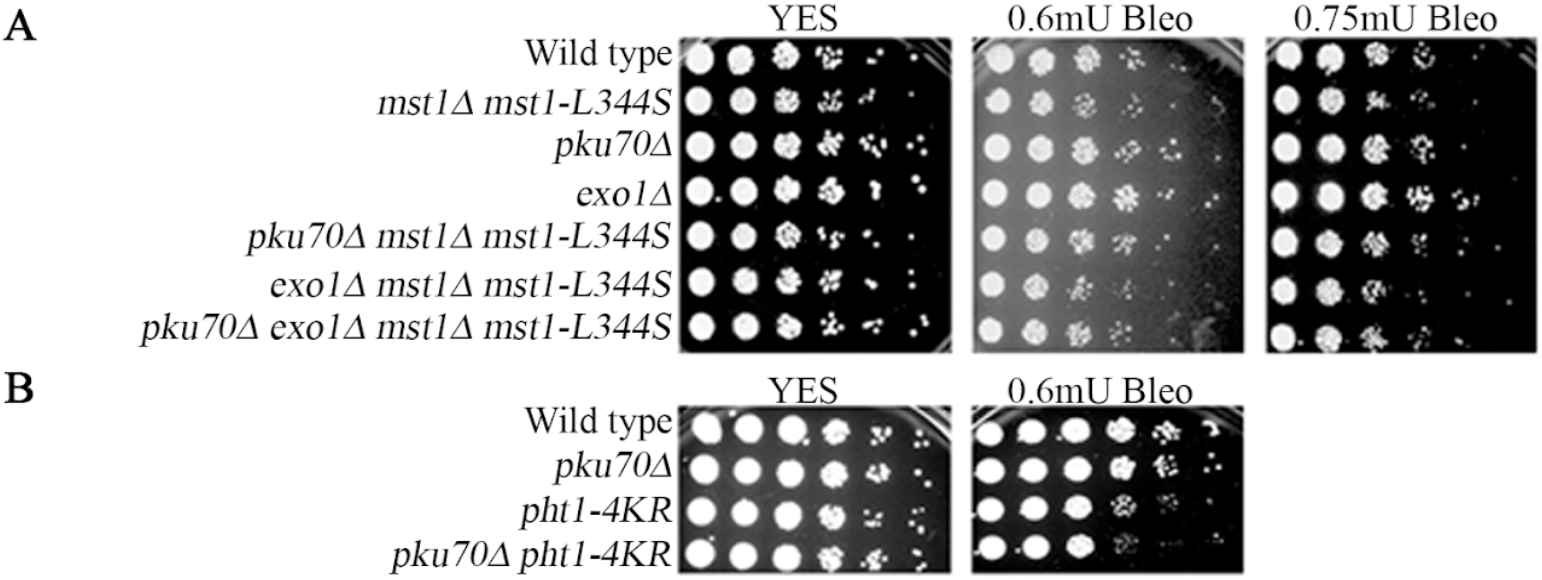
*mst1-L344S* mutants are sensitive to the DSB-inducing agent bleomycin. *pku70Δ, exo1Δ* **A**. and *pht1-4KR* **B**. growth in bleomycin. Cells were grown in YES media overnight at 25°C then 5X serial dilutions were spotted onto YES plates or YES plates containing bleomycin. Plates were incubated at 25°C and photographed after 4 days. 1 unit of bleomycin = 1 mg (by potency)

We wondered whether Pku70 also affects the recruitment of repair proteins in *mst1-L344S*, so we imaged fluorescently tagged RPA under phleomycin treatment. We observed first that the percentage of untreated cells with RPA foci in *pku70Δ* and *pku70Δ mst1-L344S* is higher than in wild type or *mst1-L344S* (Figure 2A, supplementary figure 2, supplementary table 1), suggesting a burden of endogenous damage. However, cell growth and cell length at septation were not significantly affected (Figure 3A, supplementary figure 3).

Upon treatment, the number of positive cells is only slight increased, and does not reach the level observed in wild-type cells (Figure 2A, supplementary figure 2A, supplementary table 1). Results were slightly different for Rad52: the *pku70Δ* single mutant cells showed a comparable level to wild type in untreated conditions, with little change following treatment. The untreated double mutant *pku70Δ mst1-L344S* were higher than either single mutant and showed a modest increase upon treatment, but not to the levels of wild type (Figure 2A, supplementary figure 2A, supplementary table 1). Finally, the fraction of *pku70Δ* with Rad54 foci is similar to wild type in untreated cells, increased by the same relative proportion following treatment. Again, the untreated *mst1-L344S pku70Δ* has a higher baseline in untreated cells than either single mutant, and induces to approximately wild type levels (Figure 2B, supplementary figure 2C, supplementary table 1).

We used the mCherry-LacI marked HO-induced DSB system to examine the apparent resection of DSB in *pku70Δ* mutants by monitoring the disappearance of mCherry signal (Figure 2D). We find that *pku70Δ* single mutant cells do show a slight reduction in mCherry-DSB foci over time, but this effect is far more modest than that in wild-type. At the end of the time-course, just under 70% of the *pku70Δ* cells still contain mCherry-DSB foci. We observed a similar phenotype in *pku70Δ mst1-L344S* cells. (Figure 2D).

Long range resection is initiated by MRN and extended by the exonuclease Exo1, as well as other proteins (Zhu *et al*. 2008; Langerak *et al*. 2011; Yan *et al*. 2019). We observed that mCherry-LacI marked DSBs do not disappear over the time-course in *exo1Δ*, in contrast to wild-type (Figure 2D), and consistent with a failure in resection. We observed that *exo1Δ, mst1-L344S*, and the double mutant *exo1Δ mst1-L344S* all show a similar persistence of the mCherry signal. Additionally, *exo1Δ mst1-L344S* has similar sensitivity to bleomycin as the *mst1-L344S* single mutant and this sensitivity is rescued by *pku70Δ* (Figure 3A). These data suggest that *mst1*^+^ and *exo1*^+^ are in the same pathway.

### Is H2A.Z a target of Mst1 to regulate DSB repair?

In other organisms that the presence and turnover of the histone variant H2A.Z (*S. pombe* Pht1), a KAT5 substrate, is important for the proper repair of DSBs by regulating resection (Xu *et al*. 2012; Gursoy-Yuzugullu *et al*. 2015; Jiang *et al*. 2015). In human cells, H2A.Z depletion leads to higher ssDNA accumulation and lower DSB recovery from HR (Xu *et al*. 2012; Jiang *et al*. 2015). Failure to remove H2A.Z at DSBs in human cells leads to higher RPA binding and higher recovery from HR (Gursoy-Yuzugullu *et al*. 2015). In fission yeast, H2A.Z is a known substrate of Mst1, and a mutant *pht1-4KR* that disrupts the Mst1 acetylation sites causes genome instability and anaphase defects (Kim *et al*. 2009). To investigate the role of H2A.Z acetylation in repair, we compared the phenotype of *pht1-4KR* and *pht1Δ* to *mst1-L344S*. We observed significant reduction in RPA and Rad52 foci in *pht1-4KR* at 25°C similar to that observed with *mst1-L344S* cells (Figure 2A, supplementary figure 2B, supplementary table 1). However, in contrast to *mst1-L344S*, the sensitivity to bleomycin we observe in *pht1-4KR* is not rescued by *pku70Δ* (Figure 3B).

We also compared the phenotypes of *pht1Δ* and *pht1-4KR* cells. *pht1Δ* showed fewer cells with RPA foci compared to wild type under phleomycin treatment (Figure 2A, supplementary figure 2B, supplementary table 1). Rad52 foci were also reduced in *pht1Δ* compared to wild type under phleomycin treatment (Figure 2A). However, the effect of *pht1-4KR* cells was more dramatic than *pht1Δ*, and much closer to the observations in *mst1-L344S* (Figure 2A, supplementary table 1). Finally, we used the mCherry-DSB reporter system to examine resection *in vivo*. As with *mst1-L344S* and *exo1*Δ, *pht1-4KR* cells showed a persistent mCherry-DSB signal, consistent with a failure to resect (Figure 2D).

### The damage checkpoint is normal in *mst1-L344S*

Previously, we showed that the temperature-sensitive allele *mst1Δ leu1*^+^*::nmt-mst1-L344S* is sensitive to S-phase specific genotoxins including methyl methanesulfonate (MMS), which causes alkylation damage that inhibits replication, and camptothecin (CPT), a topoisomerase-I inhibitor that causes lesions during S-phase (Wan *et al*. 1999; Fung *et al*. 2002; Morishita *et al*. 2003; Frampton *et al*. 2006). In contrast to our observations with phleomycin, which induces general DSBs, we observed that the percentage of cells with RPA and Rad52 foci was higher in *mst1-L344S* cells treated with these S-phase specific genotoxins than that in wild-type (Figure 4A). This is consistent with these S-phase specific lesions inducing different repair pathways, including post-replication repair via translesion synthesis and template switching to repair MMS-induced damage, as well as nucleolytic cleavage of protein -DNA adducts to repair CPT-induced damages (rev. in (Alagoz et al. 2012; Arbel, Liefshitz, and Kupiec 2020)).

**Figure 4.**
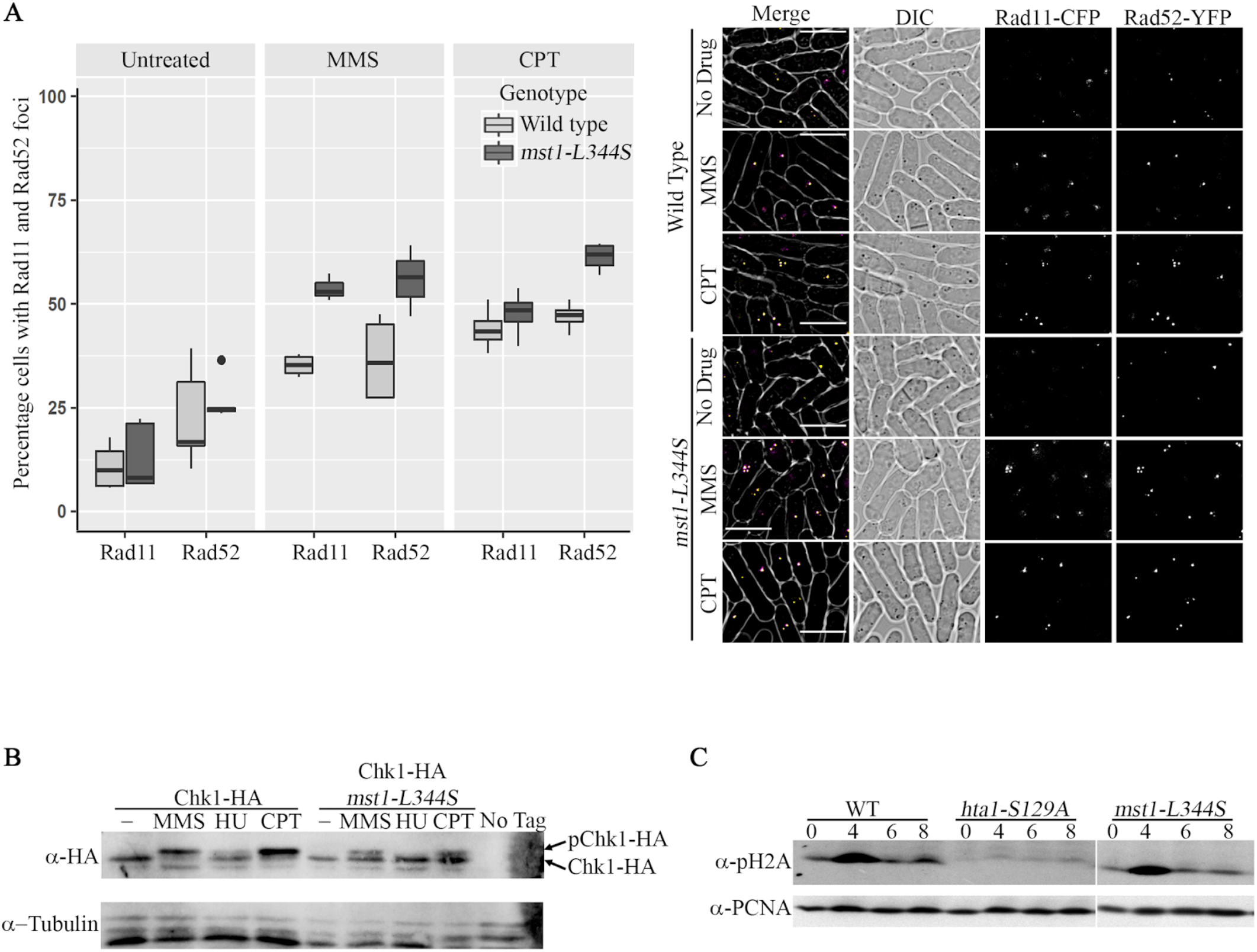
Mst1 affects DNA lesion repair but does not affect checkpoint activation. **A. Left:** Percentage of cells with RPA (Rad11) and Rad52 foci in WT and *mst1-L344S* after CPT and MMS treatments. The bold line in each box represents median and the filled circles represent outliers. **Right:** examples of WT and *mst1-L344S* cells with RPA and Rad52 foci after drug treatment. Scale bar is 10µm. Rad11-CFP is colored magenta for visibility. **B**. Phosphorylation of Chk1-HA is not affected in *mst1-L344S* after treatment of cells with three types of DNA damaging agents at 32°C. cells were treated for 4 hours with 0.01% MMS, 15mM HU or 40μM CPT and the and the Chk1-HA phosphorylation shift band was detected by Western blotting with 12CA5 anti-HA antibody. **C**. H2A phosphorylation dynamics are not affected in *mst1-L344S* when cells are treated with 40μM CPT. Western Blot with an anti-phosphorylated H2A antibody showing that phosphorylation of H2A peaks 4 hours after treatment with drug then the signal is down regulated 4 hours post drug removal. *hta1-S129A:* control non-phosphorylatable form of H2A; α-PCNA: control antibody.

We examined genetic interactions between *mst1-L344S* and mutants that affect checkpoint under treatment with a variety of S-phase specific genome stressors (Supplementary figure 4). A modest growth defect in untreated cells was observed for the double mutant *mst1-L344S rad3Δ* but not for others. In cells treated with UV or MMS at 25°C, we observed a striking negative interaction between *mst1-L344S* and damage checkpoint mutants *chk1Δ* or *crb2Δ* at 25°C, suggesting a requirement for an active damage checkpoint response under these treatments. This data is consistent with previous findings in *S. cerevisiae* that the cell cycle delay in NuA4 mutant depends on Rad9 (*Sp*Crb2 ortholog) after DSB induction (Javaheri *et al*. 2006). We treated cells with MMS, CPT or hydroxyurea (HU) and performed western blots against HA tagged Chk1. In wild-type, Chk1 is phosphorylated in cells treated with four-hour CPT or MMS, but not in untreated cells or cells treated with HU (Figure 4B; Walworth and Bernards 1996). We observed a largely similar phosphorylation pattern in *mst1-L344S* mutant (Figure 4B), although there may be a modest reduction in the amount of pChk1. H2A phosphorylation, one of the first sensors of DNA damage and double strand breaks, was also present in *mst1-L344S* cells after 4-hour CPT treatment as in wild-type cells and diminished after removing drugs from cell culture (Figure 4C). Mst1 is not required for checkpoint activation in response to S-phase specific lesions but depends on that response for viability.

## Discussion

The MYST protein Mst1 (hTIP60; ScEsa1; KAT5) is the catalytic subunit of the NuA4 histone acetyltransferase complex, and makes pleiotropic contributions to genome stability (rev. in (Lafon *et al*. 2007; Pillus 2008)). Multiple studies indicate that hTIP60 and ScEsa1 contribute to DSB repair through acetylation (rev. in (Ghobashi and Kamel 2018)), and show that the acetyltransferase binds near DSB regions (Downs *et al*. 2004; Sun *et al*. 2009; Xu *et al*. 2010; Li and Wang 2017; Cheng *et al*. 2018). The budding yeast ScEsa1 acetylates ssDNA binding protein RPA at DSBs (Cheng *et al*. 2018) and in mammalian cells, TIP60 promotes HR localization of recombination protein Rad51 (Murr *et al*. 2005).

Previously we showed that Mst1 is *S. pombe* is essential for viability and we demonstrated the temperature sensitive allele *mst1-L344S* is sensitive to a number of genotoxins (Gómez et al. 2008; Gómez, Espinosa, and Forsburg 2005). Here, we confirmed that Mst1 in *S. pombe* is important for DSB repair because both temperature sensitive alleles are sensitive to a DSB-inducing environment including treatment with phleomycin or bleomycin, or constitutive HO-nuclease expression (Figure 1A, 1C, 3 and supplementary figure 1). We find that Mst1 directly localizes to DSBs, and this localization depends on H3K9^me^ (Figure 1E and Figure 5). We observe that even under permissive growth conditions, *mst1-L344S* shifts the balance of repair away from HR towards SSA or NHEJ (Figure 1D). We also observed that a lower percentage of *mst1-L344S* cells compared to wild type show visible foci of HR associated proteins RPA, Rad52 or Rad54, following treatment with DSB inducing agents (Figure 2A, B). Together, these observations suggest that Mst1 facilitates homologous recombination to repair DSBs, and in its absence, HR is impaired (Figure 5).

**Figure 5.**
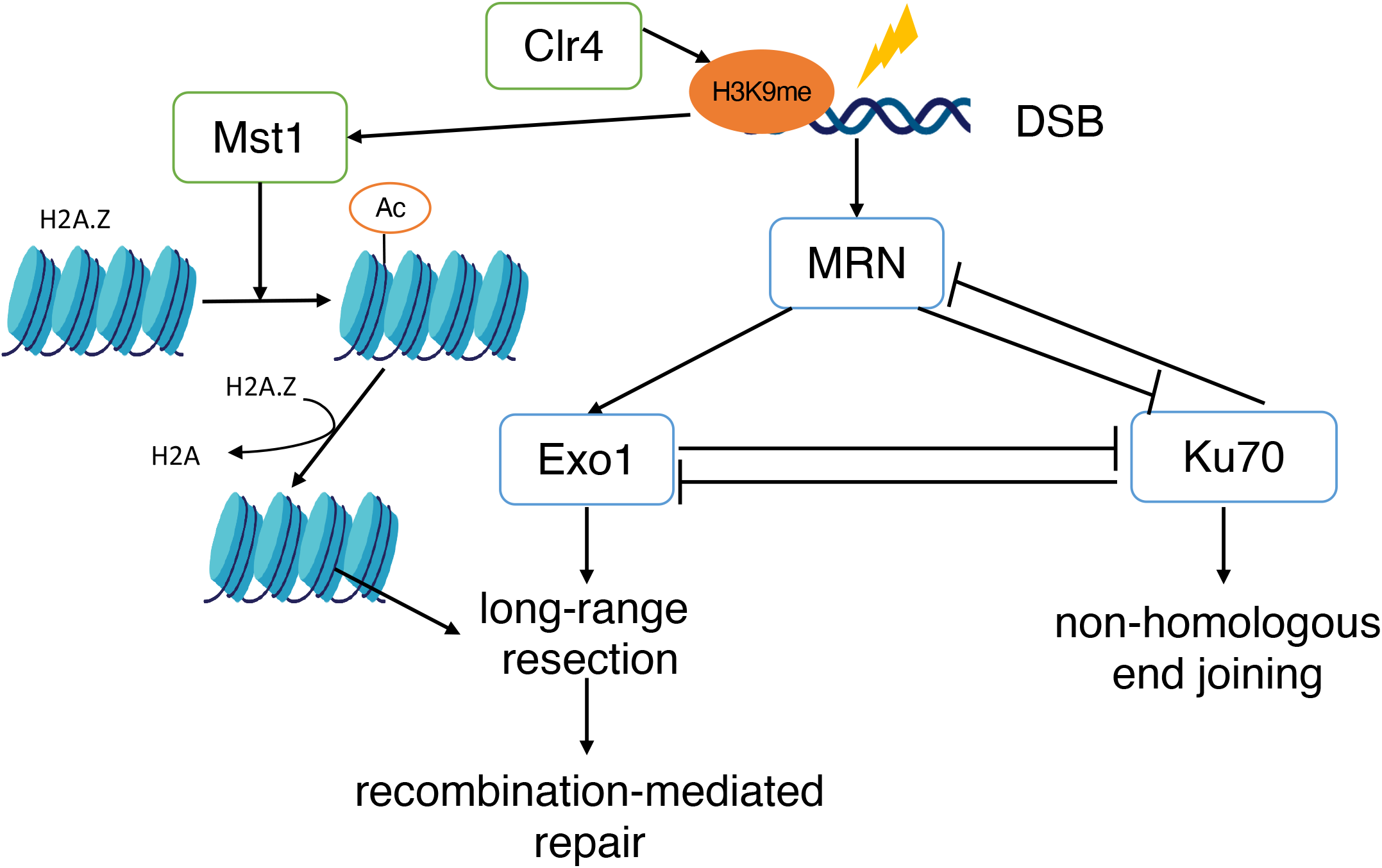
Model for the function of Mst1 in DSB repair. Mst1 facilitates HR by promoting long-range resection and inhibiting Ku70-associated mechanism that inhibits HR. Such function is accomplished by acetylating H2A.Z and thus promoting the turnover of H2A.Z at the DSB. Therefore, the histone variant can no longer block the break site. The removal of H2A.Z allows Exo1 to access the break and thus promotes downstream recombination-mediated repair.

There are conflicting data as to whether KAT5/NuA4 influences non-homologous end joining (NHEJ), which is typically the alternative pathway for repair if HR is inhibited (Chang *et al*. 2017). In budding yeast, ScEsa1 has been reported to be essential for NHEJ at DSBs (Bird *et al*. 2002). However, studies in mammalian cells suggest TIP60 instead inhibits NHEJ by acetylating H2A and H4; this reduces loading of 53BP1 protein that promotes NHEJ (Tang *et al*. 2013; Renaud *et al*. 2015; Jacquet *et al*. 2016). Consistent with this, in budding yeast deletion of the 53BP1 orthologue Sc*rad9* increases resection efficiency (Bonetti *et al*. 2015) and similarly, fission yeast *crb2*Δ mutants have increased resection (Leland *et al*. 2018). However, in fission yeast, the G2-dominant cell cycle means that NHEJ is generally not the preferred pathway for DSB repair in growing cells (rev. in (Langerak and Russell 2011)). Because of this, we did not specifically examine efficiency of NHEJ. Our data suggest that Mst1 affects homologous recombination, specifically by reducing long-range resection (Figure 2 and Figure 5).

To monitor resection, we used a visual assay in which a lacO array marked by lacI-mCherry is adjacent to an induced double strand break. Previous studies have shown that loss of the lacO array adjacent to a break correlates with resection (Yu *et al*. 2013; Leland *et al*. 2018). Consistent with this, we find that the mCherry signal declines over time in a wild type in the presence of the induced break, but not in an *exo1Δ* mutant that lacks the long-range exonuclease. We also observe that *mst1-L344S* resembles *exo1Δ* in this assay. The data from this, as well as the mutant sensitivity to DSB-inducing drugs, suggest that *mst1* and *exo1* are in a common epistasis group (Figure 5).

The end-binding protein complex Ku inhibits resection and thus, homologous recombination (rev. in (Shibata, Jeggo, and Löbrich 2018)). Its activity is coordinated with the Mre11-Rad50-Nbs1 (MRN) complex associated with early steps in resection (rev. in (Shibata, Jeggo, and Löbrich 2018; Syed and Tainer 2018)). While Ku inhibits resection driven by MRN, MRN removes Ku to facilitate homologous recombination (HR) over NHEJ (Langerak and Russell 2011; Shao *et al*. 2012; Myler *et al*. 2017; Shibata *et al*. 2018). Previous work has shown that *pku70Δ* rescues some resection in the absence of the MRN complex (Tomita *et al*. 2003; Langerak *et al*. 2011; Teixeira-Silva *et al*. 2017). Consistent with these observations, we found *pku70Δ* also partially rescues *mst1* and *mst1 exo1Δ* sensitivity to DSB-inducing agents, and partly restores recruitment of HR proteins and resection. This agrees with a model that Pku70 inhibits an alternative resection pathway that is independent of MRN, Mst1, or Exo1.

However, we observe an intermediate effect of Pku70 on resection in our visual assay: unexpectedly *pku70Δ* showed stabilization of the mCherry-LacI signal in both wild type and *mst1-L344S* backgrounds, whereas we expected at least the single mutant to show wild type levels of resection. Since Pku70 represses microhomology-mediated end joining (MMEJ), one possibility is that an increase of MMEJ antagonizes resection directly (Decottignies 2007; Wang and Xu 2017). This may also distinguish a role for Ku in the initiation of resection (which is Exo1-independent) from long range resection, which is likely required to see loss of the mCherry-LacI signal and could be antagonized by MMEJ in our assay. Ku also plays a role in telomere maintenance (Baumann and Cech 2000; Manolis *et al*. 2001), in replication fork restart (Teixeira-Silva *et al*. 2017), and in rDNA stability (Miyoshi *et al*. 2009; Sánchez and Russell 2015). It represses chromosome rearrangements (Prudden *et al*. 2003). We observed increased RPA signal in the *pku70Δ* mutant without any treatment, suggesting the loss of Pku70 causes an increased burden of ssDNA. It is possible that efficiency of resection at a single engineered break is reduced due to recruitment of resection activities to numerous other sites of genotoxic stress that are the result of Ku depletion, but this observation will require further analysis.

Multiple studies show the presence and the turnover of the histone variant H2A.Z (SpPht1) is required for DSB repair (Kalocsay *et al*. 2009; Xu *et al*. 2012; Gursoy-Yuzugullu *et al*. 2015; Jiang *et al*. 2015). In mammalian cells, the absence of H2A.Z leads to more ssDNA accumulation and reduced Pku70 localization (Xu *et al*. 2012; Gursoy-Yuzugullu *et al*. 2015). Exchange of H2A.Z is required for H4 acetylation by TIP60, which creates open chromatin (Gursoy-Yuzugullu, Ayrapetov, and Price 2015; Xu et al. 2010, 2012). Several acetyltransferases have been implicated in H2A.Z acetylation including KAT5 (Mst1) and KAT2A (Gcn5) (Kusch *et al*. 2004; Babiarz *et al*. 2006; Keogh *et al*. 2006; Millar *et al*. 2006; Zlatanova and Thakar 2008; Kim *et al*. 2009; Semer *et al*. 2019). In mammalian cells, the deposition of H2A.Z precedes its acetylation (Xu *et al*. 2012), while in Drosophila, the acetylation of H2A.Z homolog H2Av stimulates its removal after DNA damage (Kusch *et al*. 2004). Blocking TIP60 prevents both H2A.Z exchange and H4 acetylation (Morillo-huesca *et al*. 2010; Papamichos-Chronakis *et al*. 2011; Xu *et al*. 2012; Shastri *et al*. 2018).

To investigate whether H2A.Z acetylation by Mst1 influences repair, we used a non-acetylable mutant of H2A.Z, *pht1-4KR*. This mutation has previously been shown to disrupt Mst1 acetylation sites and leads to chromosome segregation errors (Kim *et al*. 2009). We saw a similar percentage of *pht1-4KR* cells with RPA and Rad52 foci as seen in *mst1-L344S* cells in treated cells, both significantly lower than that in wild type. Additionally, mCherry-DSB foci in *pht1-4KR* did not disappear across the time-course, and the percentage of mCherry-DSB persistence was similar to that in *mst1-L344S* (Figure 2D). This suggests that *pht1-4KR* and *mst1-344S* affect in a common pathway required for resection. We speculate that *pht1-4KR* once inserted cannot be efficiently removed, and interferes with H4 acetylation and chromatin decompaction. This would essentially lock the chromatin in a form refractory to long range resection. This could explain why sensitivity of *pht1-4KR* to DSBs is more severe than that of *pht1Δ*, suggesting that an immobile H2A.Z is more damaging than no H2A.Z at all. However, in contrast to *mst1, pht1-4KR* sensitivity to bleomycin is not rescued by *pku70Δ* (Figure 3B). This might suggest that H2A.Z binds to DSBs after Pku70 is removed, but that would be inconsistent with work in mammalian cells suggesting that Ku-recruitment is H2A.Z dependent (Xu *et al*. 2012). Further work will be required to resolve these observations.

Finally, we show that the temperature sensitive allele *mst1Δ leu1*^+^*::nmt:mst1-L344S* is sensitive to S phase specific genotoxins including methyl methanesulfonate (MMS) and camptothecin (CPT) even at permissive temperature (Gómez et al. 2008). This is consistent with a recent study showing that Mst1 in *S. pombe* is important for CPT-induced damge repair (Noguchi *et al*. 2019). Studies in *S. cerevisiae* showed Esa1’s importance to MMS-induced damage repair through translesion synthesis and sister chromatid recombination (Bird *et al*. 2002; House *et al*. 2014; Renaud-Young *et al*. 2015). Compromised DNA damage repair could be the result of defective damage checkpoint activation. In mammalian cells, TIP60 is known to acetylate ATM and p53 (Sun *et al*. 2005; Sykes *et al*. 2006; Tang *et al*. 2006). However, we find that the temperature sensitive allele does not affect activation of DNA damage checkpoint Chk1 or phosphorylation of H2A(X) (Figure 4B and C).

Taken together, our data suggest that SpMst1contributes to repair of a variety of DNA lesions. As in other organisms, our data show that Mst1 acetyltransferase has a particular role in double strand break repair, where it facilitates homologous recombination by promoting long range resection and antagonizing NHEJ-promoting factors such as Ku (Figure 5). Our data suggest that this depends on acetylation of H2A.Z. The conservation of function of this acetyltransferase across eukaryotes indicates the fundamental nature of chromatin modification in DNA repair transactions.

## Acknowledgement

We would like to thank Li-Lin Du, Michael Keogh, Takuro Nakagawa, Eishi Noguchi, Mathew O’Connell, and Simon Whitehall for strains. We thank Ji-Ping Yuan and Amanda Jensen for assistance and technical support. We are grateful to current members of the Forsburg lab for many helpful comments and discussions throughout the course of the study, and an anonymous reviewer for helpful suggestion. This work was supported by NIH R35 GM118109 (SLF).

## Figure Legends

**Supplementary Figure S1.**
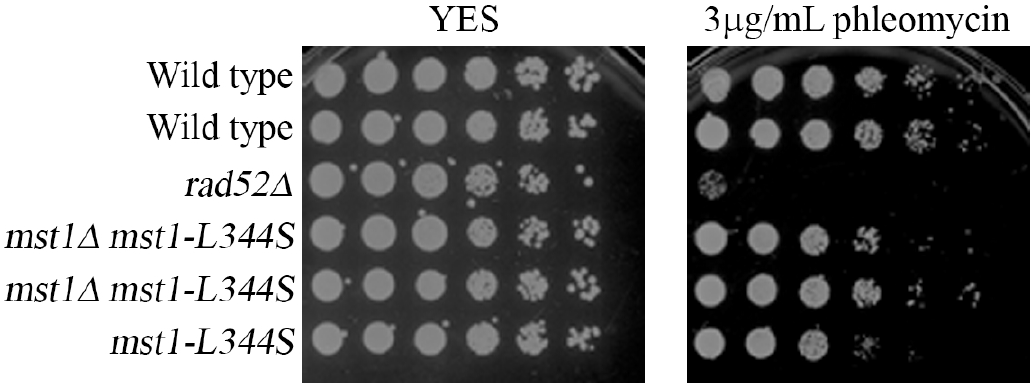
*mst1-L344S* mutants are sensitive to Bleomycin analog Phleomycin. Cells were grown in YES media overnight at 25°C then 5X serial dilutions were spotted onto YES plate or YES plates containing Phleomycin. Plates were incubated at 25°C and photographed after 4 days.

**Supplementary Figure S2.**
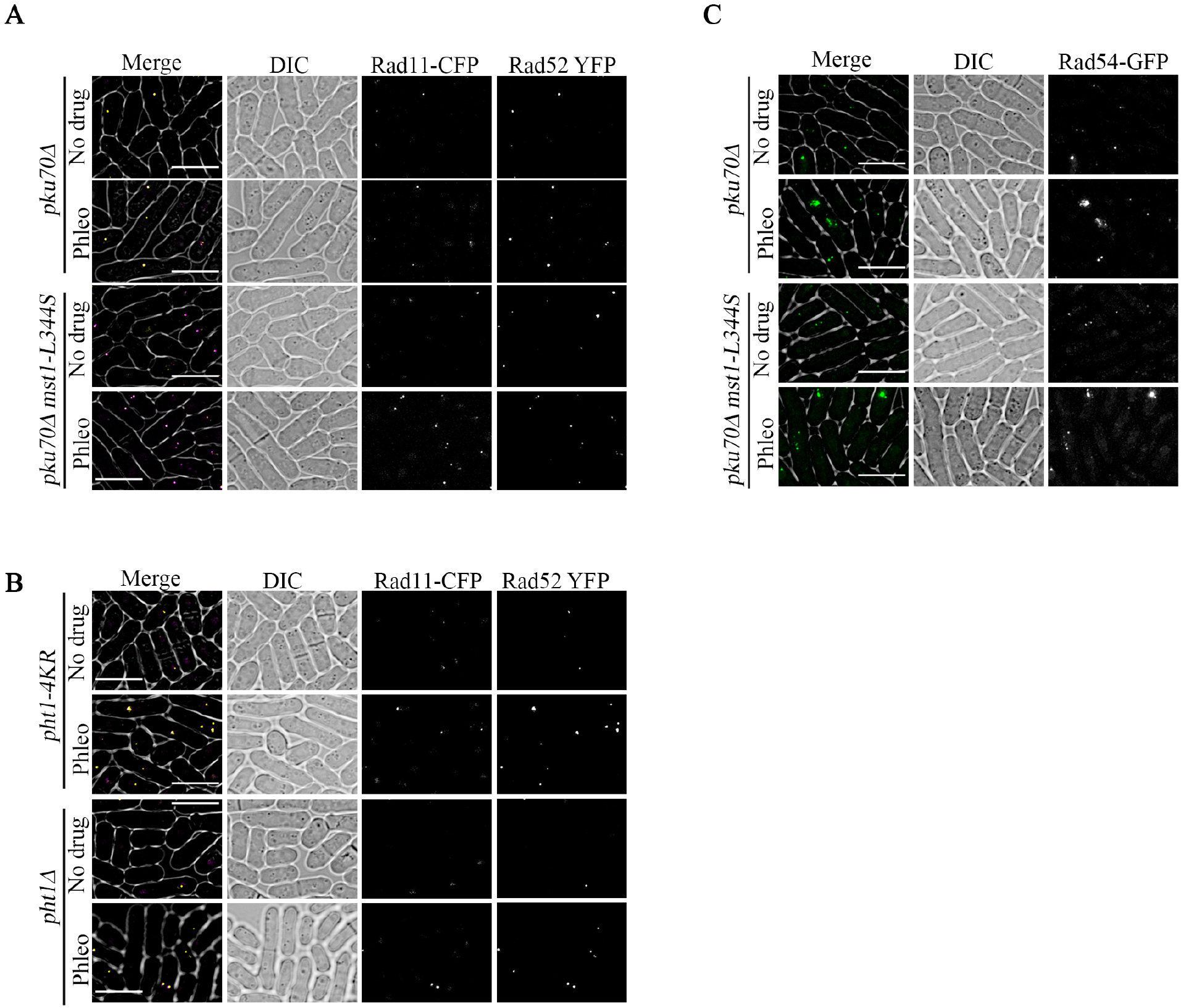
Examples of cells with RPA (Rad11), Rad52, and Rad54 foci after treatment with phleomycin at 25°C. **A**. Rad11-CFP and Rad52-YFP foci in *pku70Δ* and *pku70Δ mst1-L344S*. **B**. Rad11-CFP and Rad52-YFP foci in *pht1-4KR* and *pht1Δ*. **C**. Rad54-GFP foci in *pku70Δ* and *pku70Δ mst1-L344S*.

**Supplementary Figure S3.**
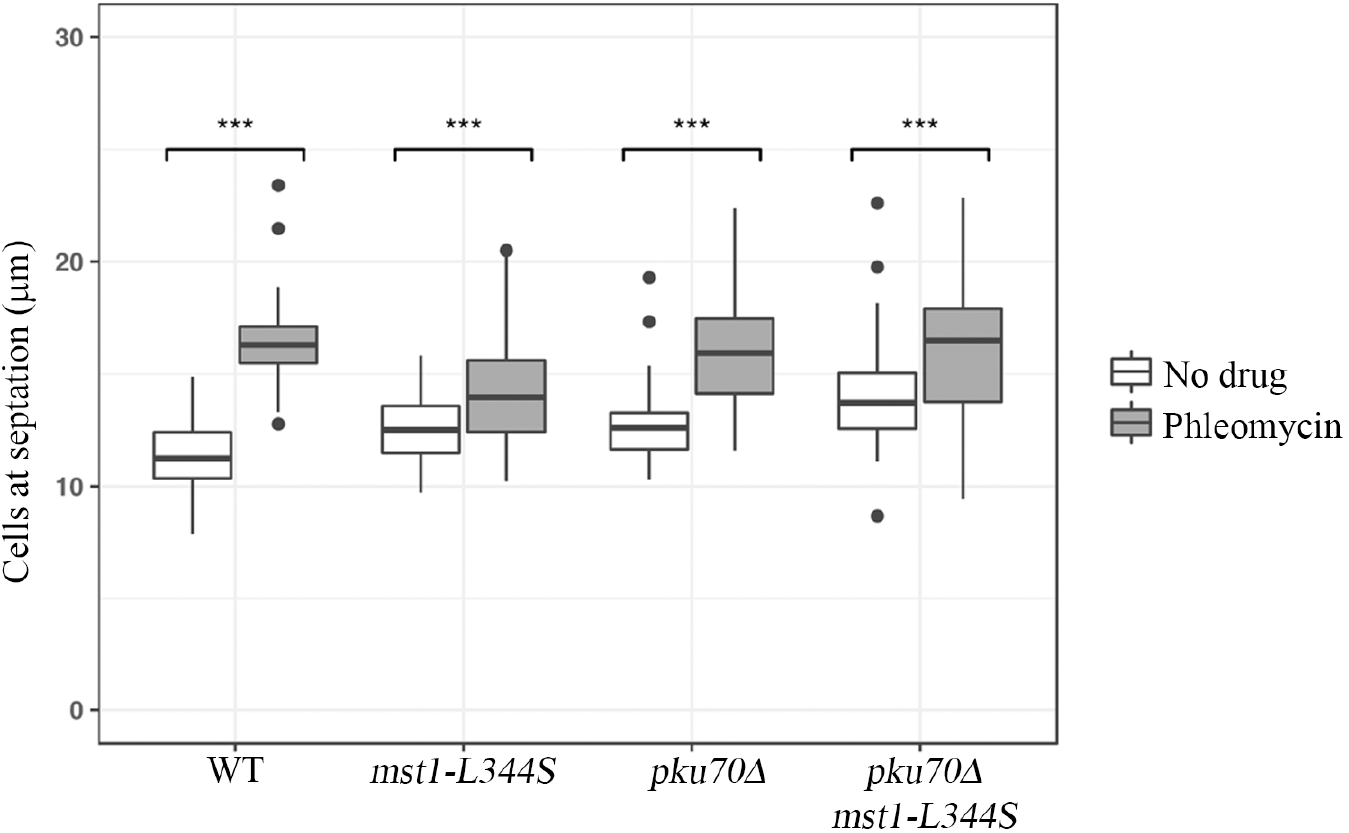
*pku70Δ* does not affect cell length at septation in *mst1-L344S* at 25°C. Length at septation at 25°C of WT, *mst1-L344S, pku70Δ*, and *pku70Δ mst1-L334S* cells. The bold line in each box represents median and the filled circles represent outliers.

**Supplementary Figure S4.**
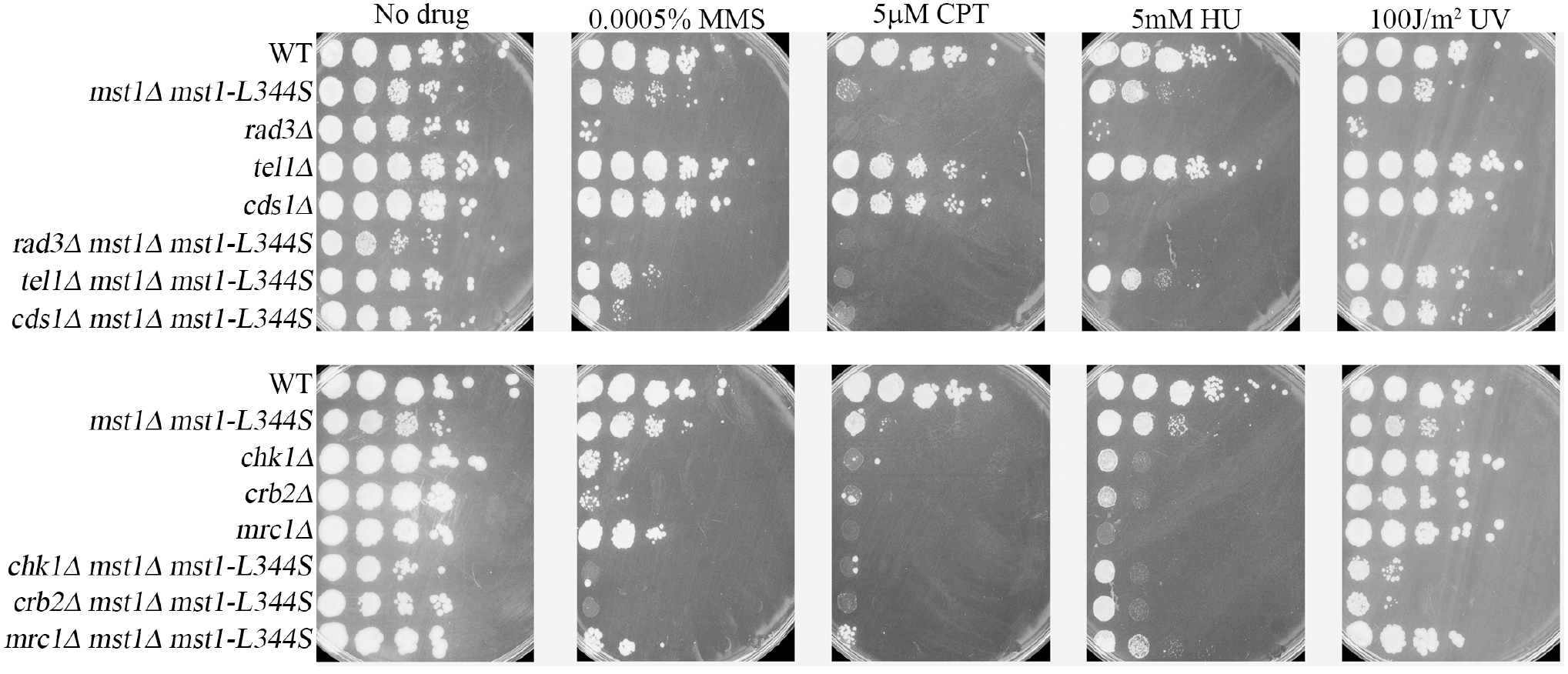
*mst1-L344S* interacts genetically with genes in the checkpoint pathways. Cells were grown in YES media overnight at 32°C then 5X serial dilutions were spotted onto YES plate or YES plates containing the indicated drugs. Plates were incubated at 32°C and photographed after 4 days.

**Supplementary Table S1.** Median, first and third quantiles of cells with Rad11, Rad52 or Rad54 percentage shown in Figure 2A and 2B.

|  | Rad11 |  |  |
| --- | --- | --- | --- |
|  | Median | 1st Quantile | 3rd Quantile |
| WT Untreated | 9.9 | 6.1 | 14.51 |
| WT Phleomycin | 43.72 | 42.98 | 50.7 |
| *mst1-L344S* Untreated | 8.14 | 6.76 | 21.15 |
| *mst1-L344S* Phleomycin | 21.3 | 15.38 | 37.29 |
| *pht1-4KR* Untreated | 3.58 | 3.25 | 4.78 |
| *pht1-4KR* Phleomycin | 24.96 | 24.76 | 25.95 |
| *pht1Δ* Untreated | 7.99 | 7.7 | 8.36 |
| *pht1Δ* Phleomycin | 39.2 | 37.75 | 39.86 |
| *pku70Δ* Untreated | 29.48 | 26.4 | 32.27 |
| *pku70Δ* Phleomycin | 38.31 | 30.99 | 46.27 |
| *pku70Δ mst1-L344S* Untreated | 29.5 | 26.38 | 31.32 |
| *pku70Δ mst1-L344S* Phleomycin | 37.08 | 36.62 | 42.3 |

|  | Rad52 |  |  |
| --- | --- | --- | --- |
|  | Median | 1st Quantile | 3rd Quantile |
| WT Untreated | 16.67 | 15.85 | 31.17 |
| WT Phleomycin | 55.4 | 53.77 | 60 |
| *mst1-L344S* Untreated | 24.73 | 24.12 | 24.89 |
| *mst1-L344S* Phleomycin | 41.85 | 35.76 | 24.89 |
| *pht1-4KR* Untreated | 11.26 | 10.98 | 11.63 |
| *pht1-4KR* Phleomycin | 36.05 | 33.84 | 38.7 |
| *pht1Δ* Untreated | 11.11 | 11.11 | 11.51 |
| *pht1Δ* Phleomycin | 43.42 | 42.53 | 45.59 |
| *pku70Δ* Untreated | 24.7 | 20.28 | 29.16 |
| *pku70Δ* Phleomycin | 27.19 | 24.71 | 31.83 |
| *pku70Δ mst1-L344S* Untreated | 35.75 | 35.2 | 42.32 |
| *pku70Δ mst1-L344S* Phleomycin | 49.09 | 46.02 | 50.4 |

|  | Rad54 |  |  |
| --- | --- | --- | --- |
|  | Median | 1st Quantile | 3rd Quantile |
| WT Untreated | 23.91 | 21.72 | 25.22 |
| WT Phleomycin | 44.91 | 37.34 | 54.55 |
| *mst1-L344S* Untreated | 22.74 | 21.18 | 23.85 |
| *mst1-L344S* Phleomycin | 32.7 | 30.28 | 47.48 |
| *pku70Δ* Untreated | 23.04 | 22.06 | 23.47 |
| *pku70Δ* Phleomycin | 47.78 | 46.09 | 49.55 |
| *pku70Δ mst1-L344S* Untreated | 30.74 | 29.99 | 31.6 |
| *pku70Δ mst1-L344S* Phleomycin | 40.97 | 40.04 | 43.37 |

**Supplementary Table S2.** Strains used in this study.

| Strain | Genotype | Source |
| --- | --- | --- |
| FY254 | *h- can1-1 leu1-32 ade6-M210 ura4-D18* | Our stock |
| FY255 | *h+ can1-1 leu1-32 ade6-M210 ura4-D18* | Our stock |
| FY527 | *h- his3-D1 ade6-M216 ura4-D18 leu1-32* | Our stock |
| FY528 | *h+ his3-D1 ade6-M210 ura4-D18 leu1-32* | Our stock |
| FY421 | *h- ade6-704 leu1-32 ura4-D18 Δchk1::ura4* | Our stock |
| FY4743 | *h- rad11-Cerulean::hphMX rad22-YFP-natMX leu1-32 ade6-M210 ura4-D18* | Our stock |
| FY8927 | *h90 mst1-L344S-5FLAG:hphMX6 rad11-Cerulean::hphMX rad22-YFP-natMX leu1-32 ura4-D18* | This study |
| FY1106 | *h+ Δrad3::ura4+ ura4-D18 leu1-32 ade6-M210* | Our stock |
| FY1204 | *h+ Δrhp51::ura4+ ade6-704 leu1-32 ura4-D18* | Our stock |
| FY1353 | *h+ leu1-32 ura4-D18 his3-D1 ade6-M210 pku70::kanr* | Our stock |
| FY2396 | *h- Δmst1::ura4+ leu1::nmt-mst1L-S leu1+ ura4-D18 ade6-M210* | Our stock |
| FY2449 | *h+ Δmst1::kanMX6 leu1::nmt-mst1L-S-leu1+ ura4-D18 ade6-M210* | Our stock |
| FY2450 | *h- Δmst1::kanMX6 leu1::nmt-mst1L-S-leu1+ ura4-D18 ade6-M210* | Our stock |
| FY2504 | *h- Δrhp9::ura4+ (=crb2) mst1::kanMX6 leu1::nmt-mst1L-S-leu1+ ura4-D18 leu1-32 ade6-M21?* | This study |
| FY2643 | *h- Δchk1::ura4+ Δmst1::kanMX6 leu1::nmt-mst1L-S-leu1+ ura4-D18 ade6-M210 or 704* | Our stock |
| FY2725 | *h- Δpku70::kanr Δmst1::ura4+ leu1::nmt-mst1L-S leu1+ ura4-D18 ade6-M210* | Our stock |
| FY2726 | *h- Δpku70::kanr Δmst1::ura4+ leu1::nmt-mst1L-S leu1+ ura4-D18 ade6-M210* | Our stock |
| FY3135 | *h- ade6-M210 leu1-32 ura4-D18 hip1::ura4+* | Simon Whitehall |
| FY3151 | *h- Δmrc1::ura4+ ura4-D18 leu1-32 his7-366* | Our stock |
| FY3439 | *h- ura4-D18 leu1-32 ade6-M216 his3-D1 Δrhp9::ura4+ (=crb2)* | Our stock |
| FY3562 | *h+ Δmst1::kanMX6 leu1::nmt-mst1L-S-leu1+ ade6-M210 ura4-D18 Δhip1::ura4+* | Our stock |
| FY3857 | *h+ Δrhp51::ura4+ Δmst1::kanMX6 leu1::nmt-mst1L-S leu1+ura4-D18 leu1-32* | This study |
| FY3884 | *h- exo1::ura4 ura4-D18* | Mathew O'Connell |
| FY4713 | *h- Δmst1::kanMX6 Δmrc1::ura4+ leu1::nmt-mst1L-S-leu1+ ura4-D18 ade6-M210 leu1-32* | This study |
| FY5304 | *h+ Δmst1::kanMX6 leu1::nmt-mst1L-S-leu1+ exo1::ura4+ ura4-D18 leu1-32* | This study |
| FY5311 | *h+ pht1-4KtoR-3HA-KanMX6 ura4-D18 leu1-32 ade6-M216 his3-D1* | Michael Keogh |
| FY6024 | *h+ mst1-L344S-5FLAG:hphMX6 leu1-32 ura4-D18* | Eishi Noguchi |
| FY6229 | *h- leu1-32 ura4-D18 ade6-M26 int::pUC8/ura4+/MATa/ade6-L469 his3-D1 ade6-M216* | Our stock |
| FY6632 | *h- pku70::kanr his3-D1 ura4-D18 leu1-32 ade6-M216* | Our stock |
| FY8929 | *h- mst1-L344S-5FLAG:hphMX6 leu1-32 ura4-D18 ade6-M26 int::pUC8/ura4+/MATa/ade6-L469 his3-D1 ade6-M216* | This study |
| FY9153 | *h+ pht1::ura4 rad11-Cerulean::hphMX rad22-YFP-natMX leu1-32 ade6-M210 ura4-D18* | Our stock |
| FY9154 | *h+ pht1-4KtoR-3HA-KanMX6 rad11-Cerulean::hphMX rad22-YFP-natMX ura4-D18 leu1-32 ade6-M216/210 his3-D1?* | This study |
| FY9338 | *h- mst1-L344S-5FLAG:hphMX6 pku70::kanr rad11-Cerulean::hphMX rad22-YFP-natMX leu1-32 ura4-D18 his3-D1? ade6-M210* | This study |
| FY9352 | *h- pku70::kanr rad11-Cerulean::hphMX rad22-YFP-natMX leu1-32 ura4-D18 his3-D1? ade6-M210* | This study |
| FY8989 | *h- rad54-GFP:hphMX6* | Takuro Nakagawa |
| FY9023 | *h- mst1-L344S-5FLAG:hphMX6 rad54-GFP:hphMX6 leu1-32? ura4-D18?* | This study |
| FY9340 | *h- pku70::kanr rad54-GFP:hphMX6 leu1-32? his3-D1? ade6-M210?* | This study |
| FY9353 | *h- mst1-L344S-5FLAG:hphMX6 pku70::kanr rad54-GFP:hphMX6 leu1-32? his3-D1? ade6-M210?* | This study |
| FY7479 | *h- arg3::Hosite-natMX ars1::p41nmt1-HO(his3+) lys1-131::pdis1-mCherry-lacI(lys1+) erg7ter::lacO(ura4+) rad52-tandemCFP::kanMX leu1-32 his3-D1 ura4-294* | Li-Lin Du |
| FY8992 | *h- mst1-L344S-5FLAG:hphMX6 arg3::Hosite-natMX ars1::p41nmt1-HO(his3+) lys1-131::pdis1-mCherry-lacI(lys1+) erg7ter::lacO(ura4+) rad52-tandemCFP::kanMX leu1-32 his3-D1 ura4-294/D18?* | This study |
| FY4620 | *h- chk1HA(ep) cds1-myc::KanMX ade6-M216 ura4-D18 leu1-32* | Our stock |
| FY9495 | *h+ exo1Δ::hph arg3::Hosite-natMX ars1::p41nmt1-HO(his3+) lys1-131::pdis1-mCherry-lacI(lys1+) erg7ter::lacO(ura4+) rad52-tandemCFP::kanMX leu1-32 his3-D1 ura4-294* | This study |
| FY9499 | *h- pku70::kanr arg3::Hosite-natMX ars1::p41nmt1-HO(his3+) lys1-131::pdis1-mCherry-lacI(lys1+) erg7ter::lacO(ura4+) rad52-tandemCFP::kanMX leu1-32 his3-D1 ura4-294/D18 ade6-M210?* | This study |
| FY9502 | *h- pht1-4KtoR-3HA-KanMX6 arg3::Hosite-natMX ars1::p41nmt1-HO(his3+) lys1-131::pdis1-mCherry-lacI(lys1+) erg7ter::lacO(ura4+) rad52-tandemCFP::kanMX leu1-32 his3-D1 ura4-294/D18?* | This study |
| FY9514 | *h+ exo1Δ::hph mst1-L344S-5FLAG:hphMX6 arg3::Hosite-natMX ars1::p41nmt1-HO(his3+) lys1-131::pdis1-mCherry-lacI(lys1+) erg7ter::lacO(ura4+) rad52-tandemCFP::kanMX leu1-32 his3-D1 ura4-294* | This study |
| FY9542 | *h- pku70::kanr mst1-L344S-5FLAG:hphMX6 arg3::Hosite-natMX ars1::p41nmt1-HO(his3+) lys1-131::pdis1-mCherry-lacI(lys1+) erg7ter::lacO(ura4+) rad52-tandemCFP::kanMX leu1-32 his3-D1 ura4-294/D18 ade6-M210?* | This study |
| FY5083 | *h- Δmst1::kanMX6 leu1::nmt-mst1L-S-leu1+ chk1HA(ep) cds1-myc::KanMX ura4-D18 ade6-M210/216?* | This study |
| FY3397 | *h- leu1-32 ura4-D18 ade6-M210 his3-D1 hta1-S129A:ura4+ hta2-S128A:his3+* | Takuro Nakagawa |
| FY5548 | *h- ura4-D18 leu1-32 his3-D1 ade6-M210 mst1::[mst1-V5 leu1+] rad22-YFP::hphMX arg3::Hosite(KanMX4) ars1::nmt41-HOendonuclease:ampR:his3+:ars1* | Our stock |
| FY9561 | *h- clr4::kanMX6-Bioneer mst1::[mst1-V5 leu1+] rad22-YFP::hphMX arg3::Hosite(KanMX4) ars1::nmt41-HOendonuclease:ampR:his3+:ars1 his3-D1 ura4-D18 leu1-32 ade6-M210* | This study |

